# Multimodal longitudinal diffusion-MRI analysis of the neocortex in a rat model of cortical dysplasia

**DOI:** 10.1101/2024.07.09.602800

**Authors:** Paulina J. Villaseñor, Hiram Luna-Munguía, Alonso Ramirez-Manzanares, Ricardo Coronado-Leija, Luis Concha

## Abstract

The neocortex is a highly organized structure, with region-specific spatial patterns of cells and fibers constituting cyto- and myelo-architecture, respectively. These architectural features are modulated during neurodevelopment, aging, and disease. While invasive techniques have contributed significantly to our understanding of cortical patterning, the task remains challenging through non-invasive methods. Structural magnetic resonance imaging (MRI) has advanced to improve sensitivity in identifying cortical features, yet most methods focus on capturing macrostructural characteristics, often overlooking critical microscale components. Diffusion-weighted MRI (dMRI) offers an opportunity to extract quantitative information reflecting microstructural changes. Here we investigate how different dMRI modalities contribute to the detection of microstructural characteristics and whether per-bundle approaches can disentangle characteristics related to the orientational organization of the myelo- and cyto-architecture in an animal model of cortical dysplasia, a malformation of cortical development. We scanned 32 animals (n=16 experimental; n=16 control) at four different time points (30, 60, 120, and 150 post-natal days) using both structural and multi-shell diffusion-weighted MRI. All dMRI metrics were sampled using a 2D curvilinear system of coordinates as a common anatomical descriptor across animals. Per-bundle metrics were labeled according to their orientation with respect to the cortical surface, and analyzed separately. Experimental animals showed diffusion abnormalities of the tangential and radial fiber components in deeper cortical areas, consistent with histological findings of neuronal and fiber disorganization. The ability of dMRI to detect abnormalities in an animal model of cortical dysplasia is indicative of the clinical potential of advanced dMRI methods to study cortical microstructure in neurological disorders.

## 1 Introduction

The architecture of the human cerebral cortex is complex, yet stereotypical. Neurons and glial cells are precisely arranged in horizontal layers, with vertically-aligned neurons forming microcolumns [25, 56, 66]. A parallel cortical organization, termed myeloarchitecture, is created by the neuropil, with axons and dendrites running radially and tangentially to the cortical surface, supporting inter-columnar communication, as well as input and output to and from the cortex. Decades ago, Vogt & Vogt observed a consistent spatial arrangement of myelin across the cortex, albeit with variations in density and fiber length according to each cortical area [61, 78]. These spatially-varying histological patterns are complementary to the cytoarchitectonic areas described by Brodmann, and are related to functional roles [18]. Historically, the identification of cyto- and myelo-architectural patterns throughout the human neocortex—and their deviations from normality—has been relegated to invasive (and destructive) techniques. However, the examination of these patterns through non-invasive methods such as magnetic resonance imaging (MRI) would provide valuable information for the study of cortical development and aging, as well as for the identification of alterations of cortical structure in neurological disorders.

In recent decades, the field of MRI has experienced rapid progress transforming our ability to visualize and quantify cortical structures. For instance, parcellation of cortical areas based on contrast intensity from Tl or T2-weighted images enabled differentiation between cortical areas, measurement of cortical thickness, and visualization of myelination across the cortical surface [16, 30, 36]. Other advanced methods such as high-resolution MRI have revealed laminar patterning features associated with myelin and cellular density within specific cortical areas [28, 79]. However, these methods still primarily capture structural aspects, often overlooking important microstructural characteristics at the sub-voxel level, particularly relevant for studying normal and abnormal neúrodevelopment, and cortical degeneration due to aging or neurological disorders. In this context, diffusion-weighted MRI provides a path to evaluate the cortex *in vivo* due to its ability to assess tissue properties at the micrometer scale by measuring the behavior of water displacement patterns over time. Though diffusion-weighted MRI has been widely applied in white matter structures due to its ability to assess fiber orientations and tissue characteristics [11, 19, 47]-advances related to image acquisition, pre-processing algorithms, the use of strong diffusion gradients, and novel computational methods have opened a new avenue to explore complex brain structures such as gray matter [27].

For many years the gold standard diffusion-weighted MRI technique has been diffusion tensor imaging (DTI; [9] for its reliability and relatively straightforward application. For example, some have applied DTI *in vivo* to differentiate between the motor and somatosensory cortices by their anisotropy features [53] or to discriminate between cortical areas [34, 57]. Others have investigated the myeloarchitectural laminar composition at high-resolution in the radial direction using ultra-high magnetic fields [7]. Nevertheless, it is well known that conventional DTI has its own limitations starting with the assumption that diffusion occurs in a free and unrestricted environment following a gaussian distribution of water displacement, which inevitably affects its ability to properly characterize complex microstructural configurations, such as the one in the cortex. To overcome the limitations of DTI, other diffusion-weighted MRI methods aim to achieve a greater and more comprehensive understanding of diffusion properties linked to microstructural features. An example of this is the immediate extension of DTI: diffusion kurtosis imaging (DKI) [43], which quantifies deviations of water diffusion displacement from gaussian distribution, providing a higher order description of diffusion hindrance and restriction within the tissue. Because kurtosis reflects how water molecules interact with cellular barriers, it serves as an indirect-measure to infer how “complex” or packed an environment is. For example, recent-studies have used DKI to investigate the inflammatory response such as gliosis [37, 85], and neurodegenerative processes including demyelination, axonal swelling [39, 41] and amyloidosis [64].

In recent years, there has been growing interest in evaluating tissue microstructure by modeling the diffusion weighted MRI signal according to prototypical biophysical compartments. Several biophysical models have been proposed in order to identify white matter characteristics like axonal radius, intra- and extra axonal diffusivities, and their relative volume fraction [1, 6, 62, 84] proving to be useful for the study of neurodegenerative diseases [3]. However, modeling the cortex is less straightforward due to the broader components found in gray matter (i.e. glia, neurons, and fibers). Previous studies have applied neurite orientation dispersion and density imaging [84] to investigate microstructural variations in the cortex [32, 33], changes in the neurite dispersion in chronic psychosis spectrum disorders [45, 59], and also in focal cortical dysplasia, were they reported a reduced intracellular volume [50]. Similarly, multidimensional tensor encoding has been useful for enhancing the delineation of cortical malformations by differentiating between gray and white matter boundaries [46]. New advanced methods offer further steps toward gray matter characterization by measuring soma and neurite density [63], and recognizing the important role of time-dependence and exchange diffusion [42] in the cortex. However, to capture the signal contributions for each compartment, high acquisition requirements are needed, including several diffusion sensitizations (up to 5 shells) and very high b-values (>10,000 s/mm^2^), resulting in lengthy scanning times.

On the other hand, diffusion-weighted MRI methods that are able to resolve per-bundle orientations can provide valuable insights not only into architectural features but-also into the specific orientations along which these structures are organized. Constrained spherical deconvolution (CSD; [73] can extract the apparent fiber density inferred by the orientation and distribution of the diffusion signal even in complex fiber configurations. While typically applied for the study of white matter, CSD has proven to be also adequate for investigating cortical layer variations according to their orientation [49]. Another promising approach are the multi-tensor signal representations computed with ad hoc optimization methods, such as the multi-resolution discrete-search (MRDS) [21] that can calculate up to three independent tensors for each voxel, thereby providing further information about the orientational microstructural components. In the MRDS implementation, the algorithm not only determines the optimal number of diffusion tensors to represent the underlying fiber configuration, but also independently estimates the main diffusion orientation and the diffusivity profile for each bundle. Additionally, the iterative use of dictionaries and non-linear optimization of the diffusion tensor makes MRDS robust and reliable, particularly at solving complex microstructural configurations, such as fiber crossings with individual diffusion profiles.

All dMRI methods offer unique insights into brain microstructure and provide complementary information that allows for the characterization of complex structures and the identification of microstructural changes linked to pathological, neurodevelopmental, and neuroplasticity. However, certain pathologies such as focal cortical dysplasia. (FCD) remain particularly difficult to characterize with diffusion-weighted MRI. FCD is a malformation of cortical development resulting from abnormal neuronal proliferation and differentiation, and is one of the leading causes of drug-resistant epilepsy. It-is characterized by disrupted cortical lamination and, in some cases, accompanied by dysmorphic neurons and balloon cells [15, 38, 44]. Nevertheless, the major challenge regarding FCD lies in its high heterogeneity and the fact that it can affect any cortical region, factors that significantly complicate diagnosis and early detection. Efforts to detect these lesions have largely focused on enhancing contrast in structural MRI [10, 13, 17, 35, 70, 75], which offers only limited access to microscale characteristics. In contrast, diffusion-weighted MRI—designed to probe tissue microstructure—remains underexplored as an approach for detecting the laminar disorganization and cellular abnormalities that define this pathology, yet-has the potential to overcome some of the limitations of conventional structural imaging. High-resolution DTI of surgical specimens have demonstrated alterations of diffusion metrics within FCD and the underlying white matter [31], and our group previously reported that the multi-tensor MRDS approach can successfully identify abnormal cortical regions in an animal model of cortical dysplasia, at early postnatal days [77]. However, little is known about whether age influences the evolution or characterization of FCD and how different diffusion-weighted MRI modalities might detect distinct microstructural components of this pathology. To this end, this study aims to: 1) compare different diffusion-weighted MRI modalities for detecting microstructural abnormalities associated with cortical dysplasia, 2) investigate whether per-bundle methods have the ability to identify cortical orientational diffusion components, and 3) evaluate how cortical disorganization evolves across age and how these diffusion-derived measures relate to histological samples.

## 2 Methods

### 2.1 Animal Model

All procedures were performed according to protocols approved by the Institute of Neurobiology UNAM ethics review board (file 111-A) and were carried out according to the Mexican Federal regulatory laws for animal experimentation (NOM-602-ZOO-1999). All authors complied with the ARRIVE guidelines.

To induce cortical lesions in our animal model similar to the ones seen in human cortical dysplasia [58], we used an antineoplastic chemical agent 1,3-Bis (2-chloroethyl) -1-nitrosourea (BCNU, also known as carmustine) that interferes with the replication of DNA, provoking mutagenesis on the neuronal proliferation during cortical neurodevelopment [12, 81]. We injected six Sprague-Dawley pregnant rats intraperitoneally with either BCNU (20 mg/kg; n=3) or saline solution for control (n=3) at embryonic day 15 (E15). Their offspring are the animals studied here (16 rats born to BCNU-treated dams and 16 born to saline-injected control dams), and were housed with their mothers until weaning in a room with a 12 h light/dark cycle with ad libitum access to food and water. Afterward, the animals were scanned at 30, 60, 120, and 150 postnatal days with our MRI protocols and processed for histology. BCNU-treated rats did not show any gross phenotypic or behavioral abnormalities compared to the controls and did not present spontaneous seizures, as expected in this animal model [4, 12].

### 2.2 *In vivo* MRI acquisitions

Acquisition protocols were carried out at the National Laboratory of Magnetic Resonance Imaging using a 7 T Bruker animal scanner equipped with a 2×2 head coil and gradients with a maximum amplitude of 760 mT/m. Animals were first anesthetized with a mix of isoflurane (4% for induction and *‘2%* for maintenance) and oxygen (2.5 1/min) during the entire scanning time while temperature and vital signs were monitored. We acquired anatomical T2-weighted images to assess the cortical morphology and diffusion-weighted images to describe the microstructure. Both sets of images were acquired in an orientation parallel to the coronal suture. Structural images were obtained using a T2-weighted Turbo-RARE sequence with the following acquisition parameters: TR/TE= 4212.78/ 33 ms and a spatial resolution of 0.117 x 0.117 x 1 mm^3^ (0.2 mm inter-slice gap). The acquisition time was 5 minutes. Diffusion-weighted images were acquired with pulsed gradient diffusion sensitization in 90 different directions, each one with b values of 600, 1200, and 2000 s/mm^2^, with diffusion gradient duration of δ=3 ms and separation Δ=9 ms. Additional fourteen b=0 s/mm^2^ images were also acquired. The sequence used an echo-planar (EPI) readout-with the following parameters TR/TE=2000/22.86 ms and spatial resolution of 0.175 x 0.175 x 1.0 mm^3^ (0.25 mm inter-slice gap). The scan time was 19 minutes.

### 2.3 Diffusion-MRI analysis

Diffusion-weighted images were first-pre-processed using tools available in MR-trix3 (ver. 3.0.3) and FSL (ver. 6.0.2) by: 1) reducing signal noise [76], and 2) the distortions induced by eddy currents [2], and 3) correcting bias field [74].

Once images were pre-processed, for each animal we manually delineated the superior (pial surface) and the inferior (intersection between the gray and white matter) cortical borders of the right hemisphere in a single slice at the level of the dorsal hippocampus (Bregma 5.86 - 3.14 mm), using ITK-SNAP [83]. Next, we created a standardized curvilinear grid to describe the cortex by fitting a Laplacian field within the manually-defined boundaries [48] (Fig. 1A; panel iii); further details are described in [77]. Then, at each point along the curvilinear grid, we sampled metrics derived from the various dMRI methods. From DTI [9], we sampled mean diffusivity (MD), axial diffusivity (AD), radial diffusivity (RD) and fractional anisotropy (FA); maps were generated using MRtrix. DKI [43] provided mean kurtosis (MK), axial kurtosis (AK), radial kurtosis (RK) and kurtosis fractional anisotropy (KFA), computed with DIPY (Python v.3.12.4). In the case of NODDI we used DIPY to compute the orientation dispersion index (ODI), free water fraction (FWF) and neurite density index (NDI) maps.

**Figure 1:**
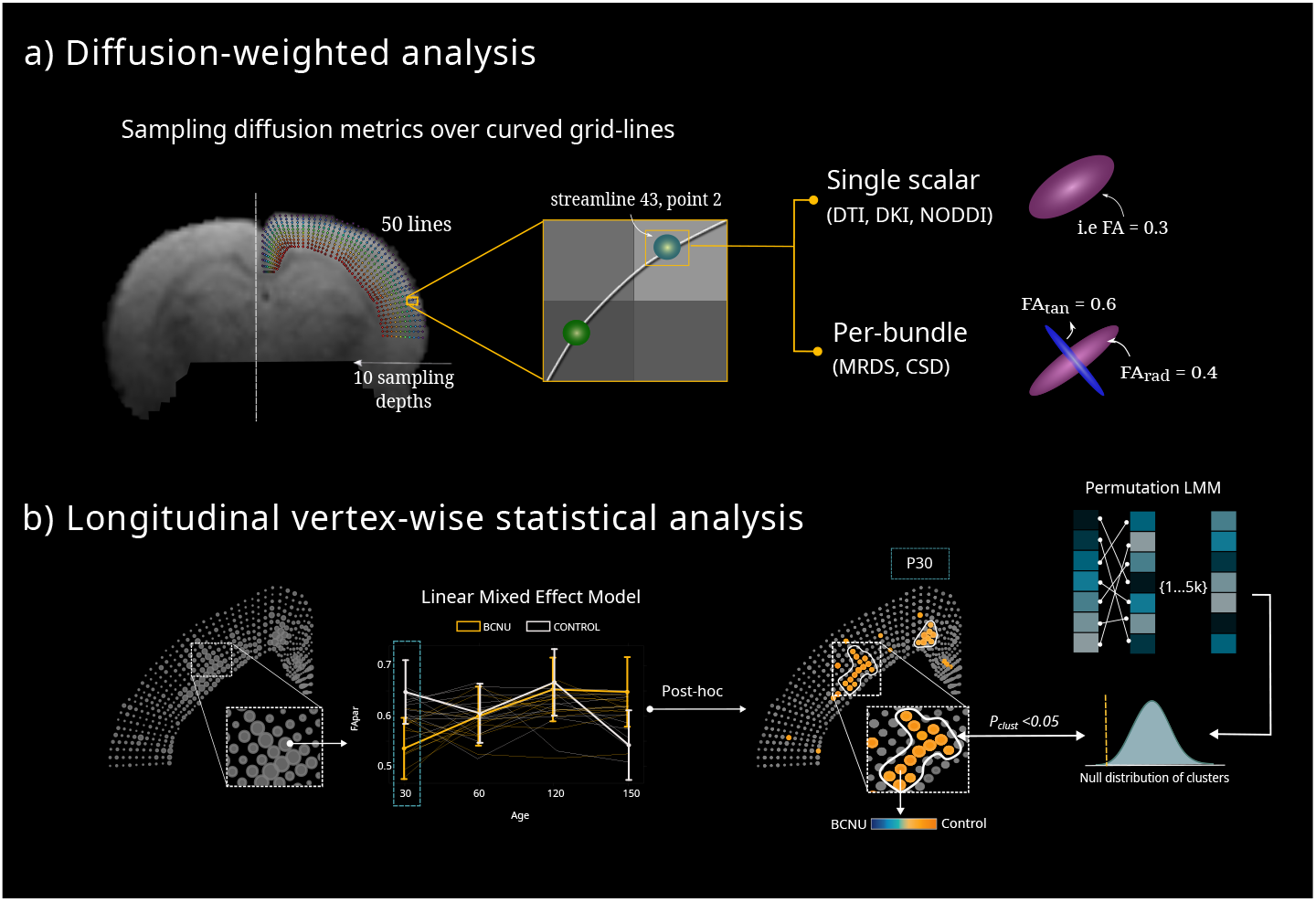
Diffusion-weighted MRI processing and statistical analysis, a) After preprocessing, quantitative maps derived from the DTI, DKI, NODDI, CSD and MRDS were sampled at each vertex. In the case of per-bundle diffusion metrics, these were separated and labeled according to their main orientation with respect to the cortical mantle (radial, magenta tensor; or tangential, blue tensor). b) A linear mixed effect model (LMM) was fit at each vertex for each diffusion metric (marker sizes in the leftmost panel indicate effect size). This was followed by post hoc testing for the significant p-value<0.05 and Cohen’s *d* to determine the direction of the estimates (blue-orange color scale). Finally, cluster-based correction for multiple comparisons was performed by permutations (white contours).

For the per-bundle assessments, we first estimated fiber orientations using CSD [73], computing the response function via the Tax calibration [72], constraining the number of fixels equal to three, and subsequently deriving apparent-density fiber maps (AFD) from the fiber orientation distribution functions (FOD) for each fiber population (fixel). For the MRDS [20] implementation, we fit up to three independent-tensors per voxel, each one with its own diffusivity profile (eigenvalues) and orientation (eigenvectors). To determine the parameters of the intra-voxel mixture of tensors, the MRDS algorithm selects the orientations from a dictionary and then applies numerical optimization techniques and a discrete search to fit-the best-eigenvalues. Next-, the Bayes’ information criterion is used to determine the multi-tensor configuration that best-explains the diffusion signal. We set-30° as the minimum angle possible between the first eigenvectors of two independent tensors. From MRDS we obtained fractional anisotropy (per-bundle FA) and mean diffusivity (per-bundle MD) for each of the identified tensors within the voxels, which were later labeled according to their main orientation with respect to the grid lines that traverse the cortical mantle. All metrics were sampled at each time point of the longitudinal analysis. For the per-bundle orientation classification-applied to both MRDS and CSD derived-metrics-we computed the dot product between the normalized principal diffusion direction of each tensor with respect to the (normalized) closest grid line segment. Here the eigenvector with the highest absolute dot product was designated as radial (*rad*; i.e. representing fibers that radiate towards the pial surface), while the eigenvector with the lowest absolute dot product was labeled as tangential (*tan;* i.e. representing fibers running tangential to the pial surface). If more than two tensors were identified at a given voxel the most perpendicular tensor or fixel with respect to the grid-lines was reported as tangential.

### 2.4 Statistical analysis

For each vertex along the grid-lines, we fit a Linear Mixed-Effects Model (LMM) analysis using R (ver. 4.1.0). The model was fit to evaluate the interaction between the BCNU and control group across age with each diffusion metric used as the response variable. Therefore, explanatory variables (age + group) were set as fixed-effects, and individual animals were modeled as random-effects. For each predictor we calculated the estimated marginal means followed by post hoc tests to identify statistical differences. Given the multiple statistical tests, we adjusted the significant p-value by permuting and recalculating the LMM test at each vertex (5,000 permutations). Then, cluster-wise statistical inference was performed by computing the empirical null distribution of cluster sizes after the 5,000 permutations with a threshold of p_clus_<0.05 with four-point connectivity (following the ideas behind AFNI’s 3dclustsim [23]. Finally, effect sizes (Cohen’s *f2* and *d)* were estimated to determine the magnitude and direction of LMM effect at each point (**Fig**. 1b).

### 2.5 *Ex vivo* assessment

#### 2.5.1 Immunofluorescence

For our histological procedure, we sacrificed two animals from each group at each time point (P30, P60, P120, and P150). After deep anesthesia with an intraperitoneal overdose of sodium pentobarbital, animals were transcardially perfused with a 0.9% NaCI solution followed by a 4% paraformaldehyde (PFA) solution (pH 7.4). Subsequently, their brains were removed and post-fixed in fresh 4% PFA solution for 24 h at 4°C. Following fixation, the brains were immorsed until sinking in 20% and 30% sucrose solution at 4 °C for 48 h. Finally, brains were frozen using dry ice and stored at -80°C. Prior to the staining process, brains were sectioned in the coronal plane (20 μm thick), at the level of the dorsal hippocampus (Bregma 5.86 - 3.14 mm) using a Leica Biosystems 3050S cryostat. The series of sections were stored in 1X phosphate buffer solution (PBS; Sigma-Aldrich) until the immunofluorescence staining. For blocking, we immersed the brain slices for 45 minutes at 4°C in 2% Bovine Serum Albumin solution (BSA; Sigma-Aldrich) and 0.3% triton X-100 (ThermoFisher) in 1X PBS. To characterize cytoarchitecture, we employed the primary antibody anti-mouse Neuronal Nuclei (NeuN; 1:350; Abcam). To assess myeloarchitecture, we performed a double immunofluorescence staining by using the following primary antibodies: anti-rabbit Myelin Basic Protein (MBP; 1:200; Sigma-Aldrich) and anti-mouse Neurofilament 200 (NF200; 1:200; Sigma-Aldrich). All primary antibodics were incubated for 24 hours at 4°C and subsequently rinsed three times with 1X PBS for 10 minutes each. Following primary antibody incubation, the corresponding secondary antibodies (Alexa Fluor) goat anti-rabbit 555 (1:500) and goat anti-mouse 488 (1:500) were incubated for 4 hours at 4C°. The slices were then rinsed three times with 1X PBS for 10 minutes each. Finally, slices were cover-slipped with microscope cover glass.

#### 2.5.2 Image processing and structure tensor analysis

Brain slices were captured using a fluorescence Apotome-Zeiss microscope equipped with 565 and 488 nm emission filters. The microscope was connected to a console integrated with the AxioVision software (ver. 4.8). Mosaics encompassing the entire slice were created by stitching individual tiles captured with a 10X objective using the MosaiX module available in AxioVision. Image processing was performed using FIJI (ver. 2.15.0). Initially, images were processed for background removal. For the channel showing fluorescence associated with MBP, we implemented structure tensor analysis using the OrientationJ plugin [65] (https://github.com/Biomedical-Imaging-Group/0rientationJ) which is designed to calculate the primary orientation for a specified number of pixels. A Gaussian kernel of 5 pm was utilized to compute the structure tensor, estimating orientation and anisotropy from a local neighborhood. The coherency parameter was then calculated, defined as the ratio between the difference and the sum of the eigenvalues. Akin to the diffusion tensor’s FA, coherency maps represent the degree of local tissue orientation, with a coherency value of 1 indicating the presence of a dominant orientation (i.e., anisotropy) in the image, while a value of 0 signifies completely incoherent orientations within the local neighborhood [24].

## 3 Results

### 3.1 Age-dependent structural changes associated with BCNU exposure

Qualitative evaluation of morphological changes between BCNU-treated animals and controls were carried out using the T2-weighted images. In **Fig**. 2, we present representative coronal sections at two anatomical levels: one at the somatosensory cortex (top row) and the second at the dorsal hippocampus (an-teroposterior -0.4 mm and -3.14 mm, respectively from Bregma; coordinates taken from the atlas rat brain of Paxinos and Watson). No evident gross morphological differences were observed between groups at P30. However, as age progressed, BCNU-treated rats exhibited notable structural alterations, including enlarged ventricles and subsequent hippocampal atrophy, where Moroni et al. [55] reported similar hippocampal lesions in adult BCNU-treated rats. In our study, these abnormalities displayed a progressive trend over time, extending to the ventral hippocampus. Interestingly, the lesions in the ventricles were not uniformly distributed among BCNU-treated rats. Each animal showed variations in severity and location, with some showing either uni-or bilateral abnormalities.

**Figure 2:**
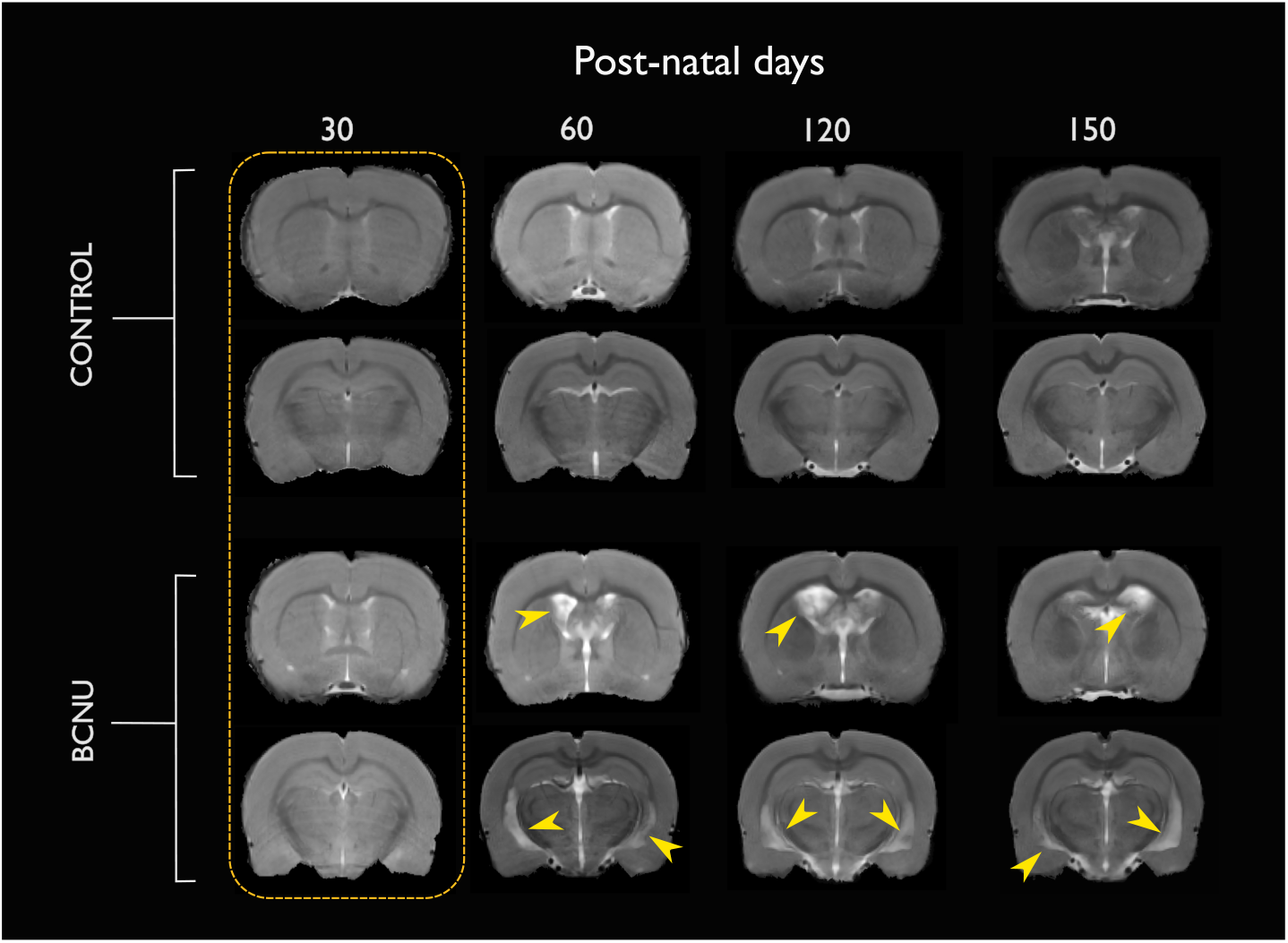
Longitudinal T2-weighted coronal images from the somatosensory cortex and dorsal hippocampus. At P30 (yellow rectangle) there were no apparent gross morphological differences between groups. BCNU-treated rats revealed progressive enlargement of the ventricles and hippocampal atrophy from P60 to P150 (yellow arrowheads). The dotted yellow box highlights lack of gross anatomical abnormalities at P30, where diffusion abnormalities were identified using dMRI (Figures 3-6).

### 3.2 Longitudinal vertex-wise diffusion-MRI analysis

#### 3.2.1 Group-average distributions

To evaluate gross group differences, we calculate the group average for each metric at each vertex along the curvilinear sampling grid-line and examine the overall distributions of values across vertices. As shown in Fig. 3, at P30 neither DTI-nor NODDI-derived metrics showed between-group differences. In contrast, DKI-derived metrics (MK, AK and RK) show more prominent group differences in the distribution of diffusional kurtosis across the grid. Although the overall shapes of the distributions are similar, the BCNU-treated animals exhibit a leftward shift in the kurtosis peak, as compared to controls. This pattern suggests a reduction in microstructural complexity or packing in the BCNU group, consistent with disruption of columnar and laminar organization. Because both AK and RK show comparable shifts, this implies that water molecules experience reduced restriction in both the radial and tangential diffusional directions. A slight increase in KFA was observed, mainly in the medial aspect-of the neocortex of the BCNU-treated animals. Overall, DKI proved to be more sensitive to microstructural changes induced by BCNU at early stages of development compared to DTI and NODDI. Longitudinal analysis **(Supplementary Fig. SI)** revealed reduced FA of superficial layers at P60 and the most medial aspect of the cortex at P150. Notably, the alterations of diffusion kurtosis tended to disappear after P30. Similarly, NODDI metrics showed variations over time, with a slight increase of NDI at P120.

**Figure 3:**
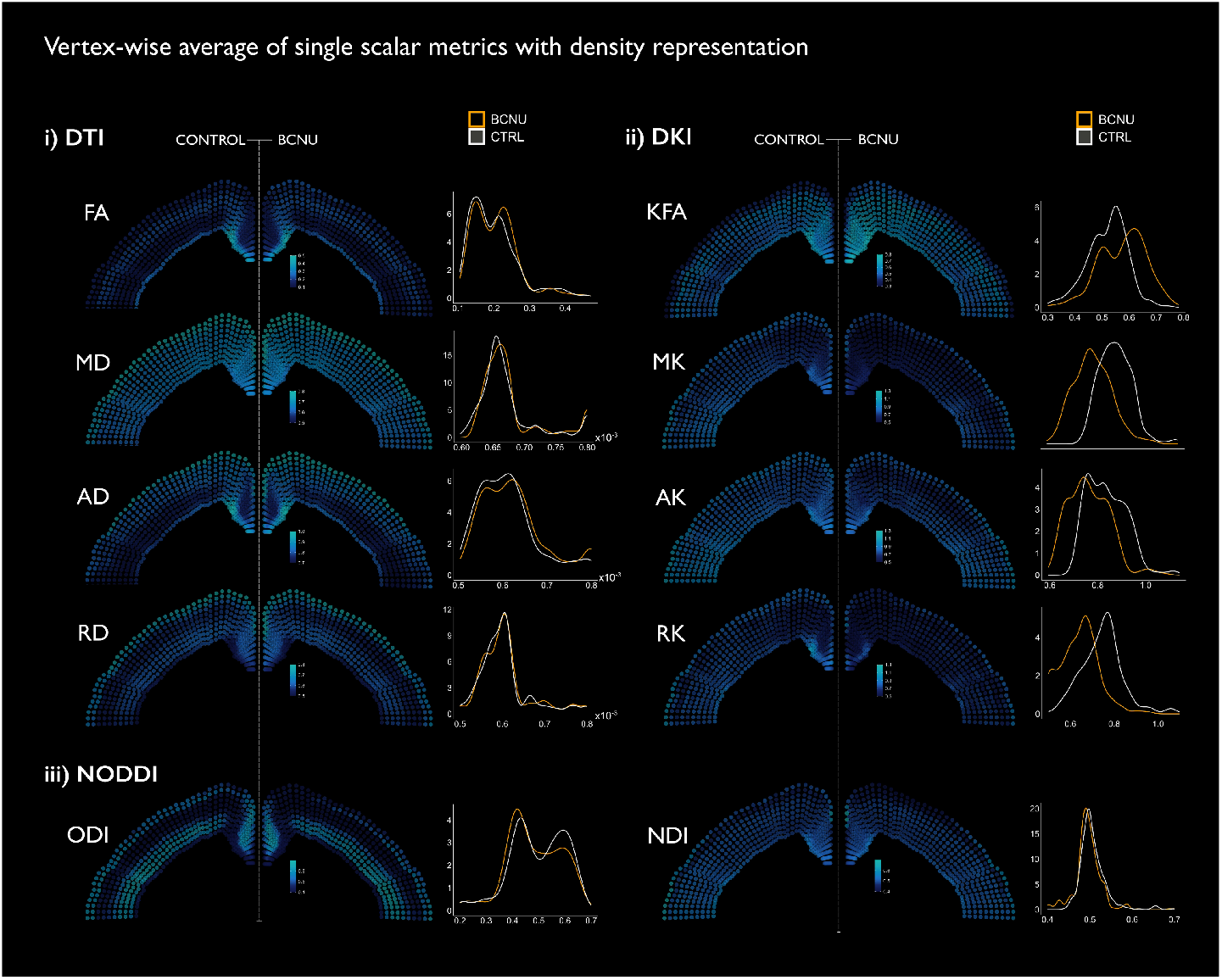
Vertex-wise means of single scalar diffusion-MRI metrics with corresponding density distributions at P30. For each vertex along the grid-line, the mean value of i) DTI, ii) DKI and iii) NODDI metrics was calculated separately for the control (hemisphere shown on the left) and BCNU (shown on the right) groups. Average values are displayed in the same blue color-coded scale for both groups. To facilitate between-group comparisons, the spatial maps for the BCNU group were mirrored horizontally. Density curves describe the frequency distribution rates of each diffusion metric across all the grid-line vertices for both groups (i.e., disregarding spatial information; BCNU in yellow, control in white).

In Fig. 4, per-bundle derived-metrics reveal consistent differences between groups across both radial and tangential diffusion orientations. In this case, FA_rad_ showed lower FA values compared to controls. This reduction is visible throughout the cortex, without clear laminar specificity, suggesting global decrease in radial microstructural coherence. This pattern is also visible at the following postnatal ages but differences between groups tended to disappear over time **(Supplementary Fig. S2)**. In contrast, FA_tan_ displays a more spatially patterned reduction. Lower FA in the tangential direction appear predominantly in the deep cortical layers, indicating a potential disruption in the tangentially-oriented cyto- and myeloarchitecture. For MD_rad_ and MD_tan_, the density plots show no major group differences between groups at any time point, suggesting that MD is less sensitive to the orientational diffusivity changes in BCNU-treated animals. The AFD measures derived from CSD align well with the FA findings. In particular, AFD_rad_ changes in BCNU-treated rats are subtle, as they show a broader and flatter density distribution, reflecting a shift towards lower radial fiber density, compared to control group, which exhibit a clearer peak distribution. This is consistent with disrupted radial organization and reduced vertical fiber density, paralleling the pattern seen in FA_rad_. Conversely, the density plot of AFD_tan_ at P30 shows a rightward shift in BCNU-treated rats, indicating higher tangential fiber density. This may reflect either an excess of horizontally oriented fibers or misoriented fibers accumulating along the tangential directions across multiple layers, which is inconsistent with normal cortical myeloarchitecture. Over time, however **(Supplementary Fig. S2)**, the increase of AFD_tan_ is not apparent. Taken together, these findings indicate a breakdown of normal radial and tangential diffusivity related to cortical columnar and laminar disorganization in the BCNU group.

**Figure 4:**
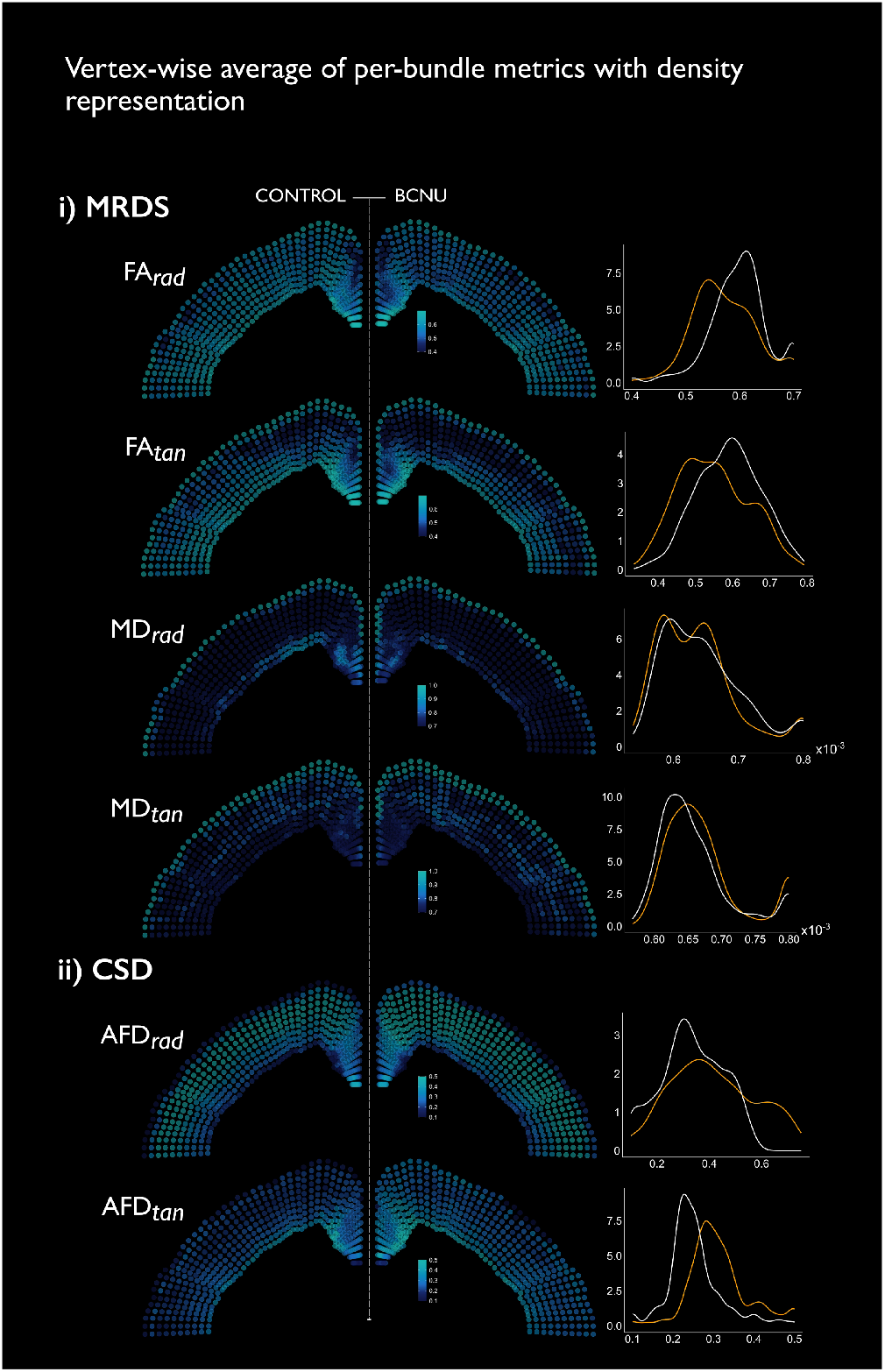
Vertex-wise means of per-bundle metrics with corresponding density distributions at P30. For each vertex along the grid-line, the mean value was computed separately for the tangential and radial directions for (i) MRDS metrics (FA, MD) and (ii) CSD apparent fiber density (AFD). Mean values are displayed using the same color scale for both groups. The right-hand panels show density estimates of the metric distributions for each group (BCNU in yellow; control in white).

#### 3.2.2 Model-based analysis

Following the group-average comparisons, we performed a linear mixed-effect model to model the longitudinal data and spatial dependency. The diffusion tensor detected only a small region of increased FA in the most superficial layers and a decrease of FA directly above the cingulum bundle in BCNU-treated rats (**Fig**. 5-i). Over time, the superficial FA reductions comprised a larger extent of the cortex at P60, but tended to disappear by P120 **(Supplementary Fig. S3)**. Mean, axial and radial diffusivities showed little (if any) abnormalities at any time point. In sharp contrast, DKI identified widespread abnormalities throughout the cortex, most notably a reduction of mean kurtosis across all cortical depths, and a reduction of axial kurtosis in the superficial layers of experimental animals at P30 (**Fig**. 5-ii). These abnormalities gradually disappeared as the animals aged, with minimal and dispersed abnormalities after P60 **(Supplementary Fig. S3)**. The spatial and temporal profiles of FA abnormalities were similar between the DTI and DKI fits. Consistent with this, NODDI revealed a focal reduction in orientation dispersion (ODI) at P30 that overlapped spatially with the region of increased FA (**Fig**. 5-iii), while NDI and FWF showed no detectable differences at this early stage. Across development, NDI displayed only minimal alterations at P60 and P120. By P150, however, control animals exhibited an increase of neurite complexity within the superficial cortical layers, whereas BCNU-treated rats failed to show this maturation **(Supplementary Fig. S4)**. At P60, an increase of ODI in BCNU animals likely indicates the presence of abnormally oriented neurites, consistent with disrupted cortical organization. The most relevant NODDI finding is at P150, where BCNU-treated rats rats demonstrated an overall decrease of FWF. This increase is a reflection of normal aging, with an expansion of the extracellular compartment, and stands in contrast to the dysplastic cortex, which retains persistent abnormal neurons and reduced laminar differentiation.

**Figure 5:**
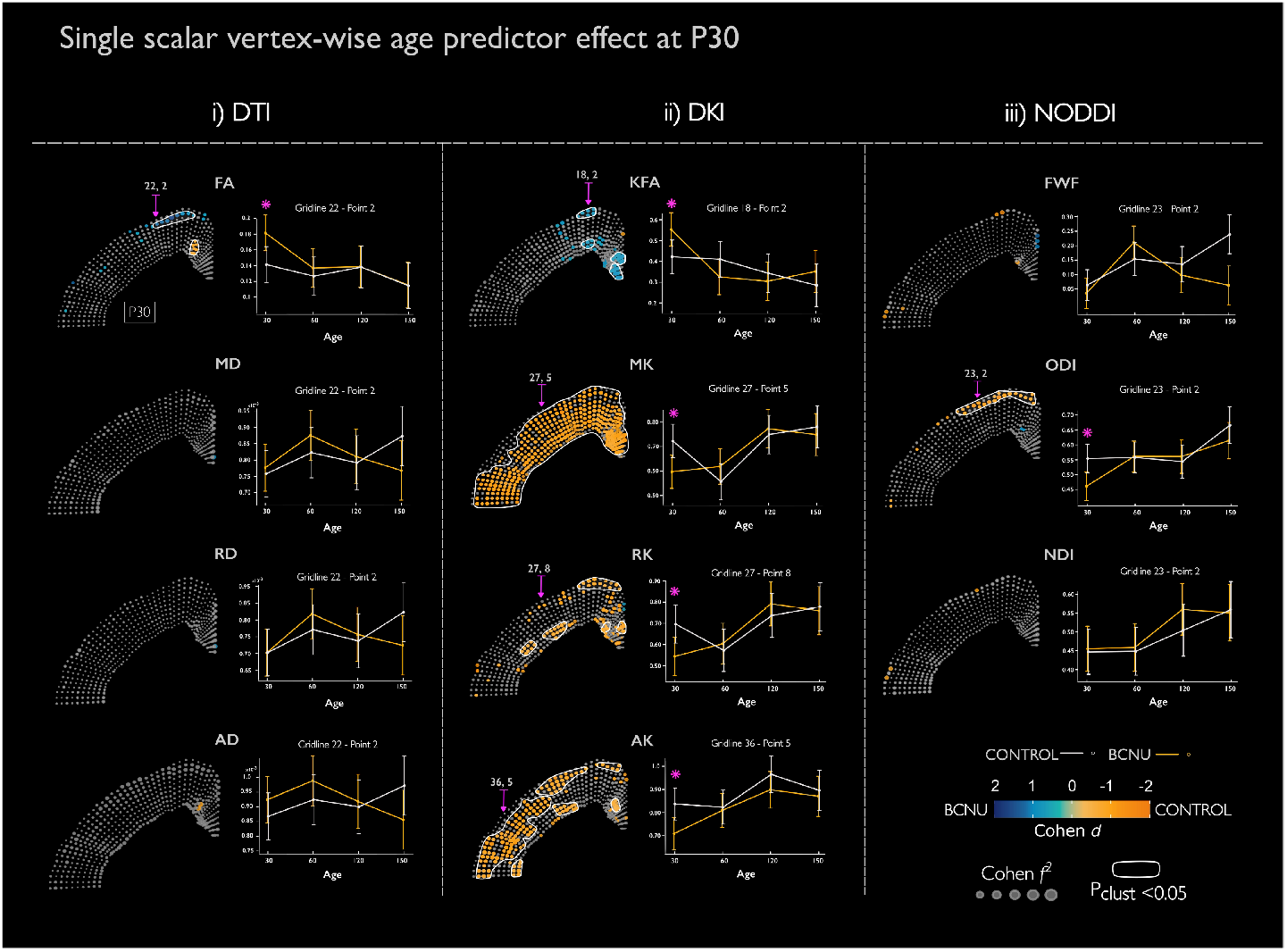
Vertex-wise analysis of the cortex at P30. Significant point-wise post hoc between-group differences (p<0.05) are represented in colors (cool colors for increases and warm colors for reductions in BCNU animals with respect to control group); non-significant in gray, with absolute effect size represented as marker size; significant clusters are represented in white contours. The LMM analyses (mean and confidence intervals) for exemplary vertices (magenta arrows) are shown to the right of each spatial map (asterisks indicate post-hoc statistical significance, p<0.05).

As compared to DTI, the multi-tensor fit performed with MRDS showed more differences between groups **(Fig. 6)**. In the ease of FA_rad_, at P30 reductions were observed at the level of the deepest cortical layers at P30. Interestingly, these changes were not evident between P60 and P120, arising again at **P150 (Supplementary Fig. S5)**, with a cluster of significant vertices located in similar cortical regions as seen at P30, but with opposite effect (i.e., abnormaily increased FA_rad_ in BCNU-treated rats). Reduced FA_tan_ was observed in deep layers at P30, but these abnormalities disappeared over time. MD_rad_ and MD_tan_ showed spread-out significant vertices across ages but without a clear spatial clustering, lacking statistical significance at the cluster level.

**Figure 6:**
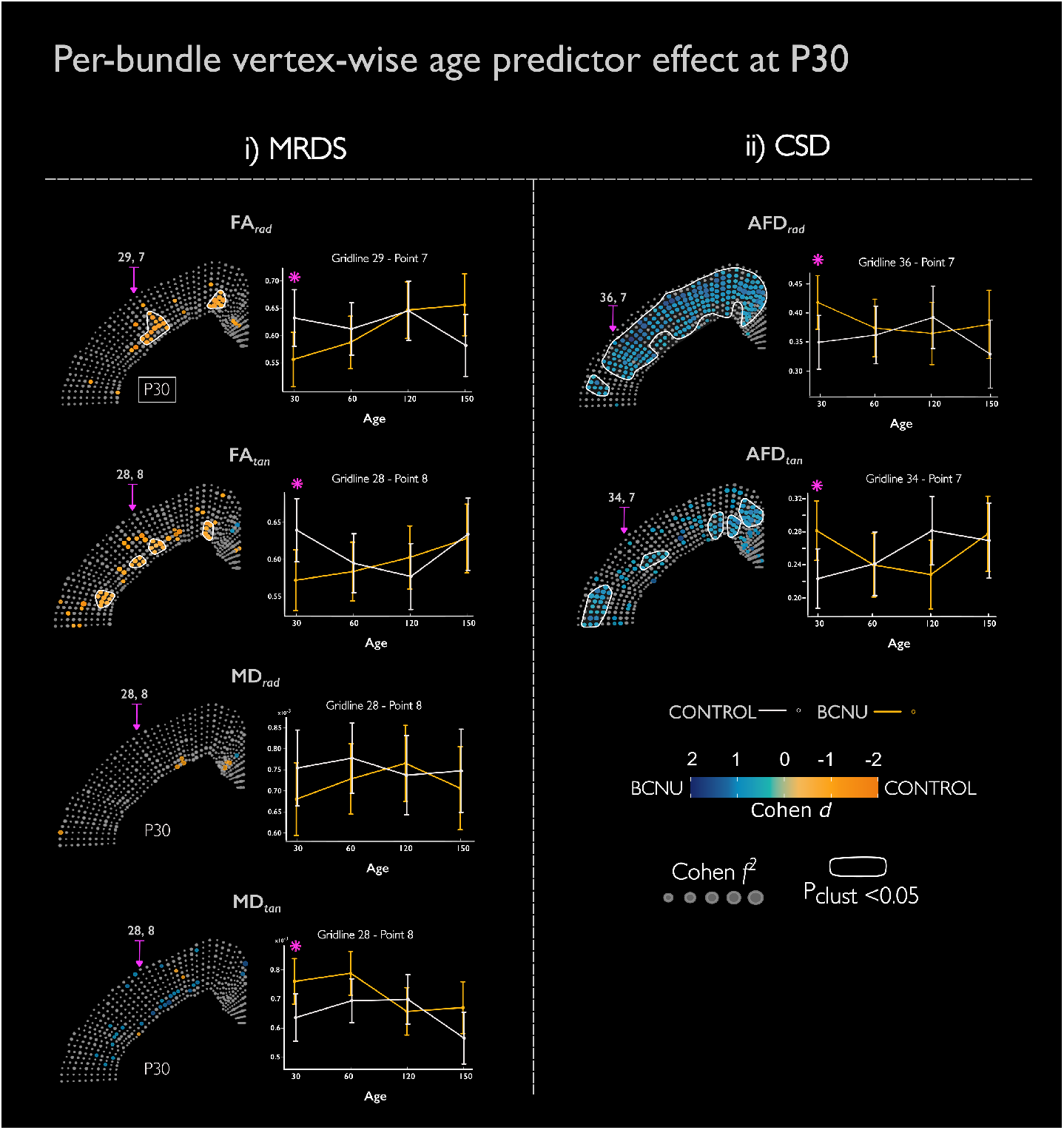
Per-bundle vertex-wise analyses with a multi-tensor fit MRDS, i), and CSD ii) at P30. Plotting conventions as in **Fig.** 5. At early stages of development (P30) FA_rad_ and FAtan, revealed significant changes (p_clus_<0.05; delineated in white) of the somatosensory and motor cortices, primarily affecting the cortical layers IV-VI. AFD_rad_ was significantly increased throughout the cortex, while AFD_tan_ was increased in smaller clusters in deeper layers. The majority of these abnormalities disappeared at later time points **(Supplementary Fig**. S6).

Per-bundle analyses performed with CSD also revealed significant abnormalities at P30. Increased AFD_rad_ was evident in a very large cluster encompassing nearly the entire cortex. Increased AFD_tan_ was also observed in several clusters situated in the middle and deepest layers. The abnormalities of both metrics, however, tended to disappear by P60 and later time points **(Supplementary Fig. S5)**

### 3.3 Histological examination

#### 3.3.1 Cytoarchitecture

The NeuN stained images yielded valuable features about the six-layered neuronal organization. Here, animals treated with BCNU showed an evident disruption of the transition between layers and their radial orientation. This abnormal pattern was evident throughout the entire brain of BCNU-treated rats, with certain cortical regions displaying more pronounced disorganization than others. Figure 7a-i presents a close-up examination of the somatosensory cortical layering from both groups. Control animals preserved a clearly defined transition across layers, with each layer showing distinct cell density, morphology, and size. On the other hand, BCNU-treated rats exhibited a blurred delineation between layers III-IV and V-VI, as indicated by the color-coded lines.

**Figure 7:**
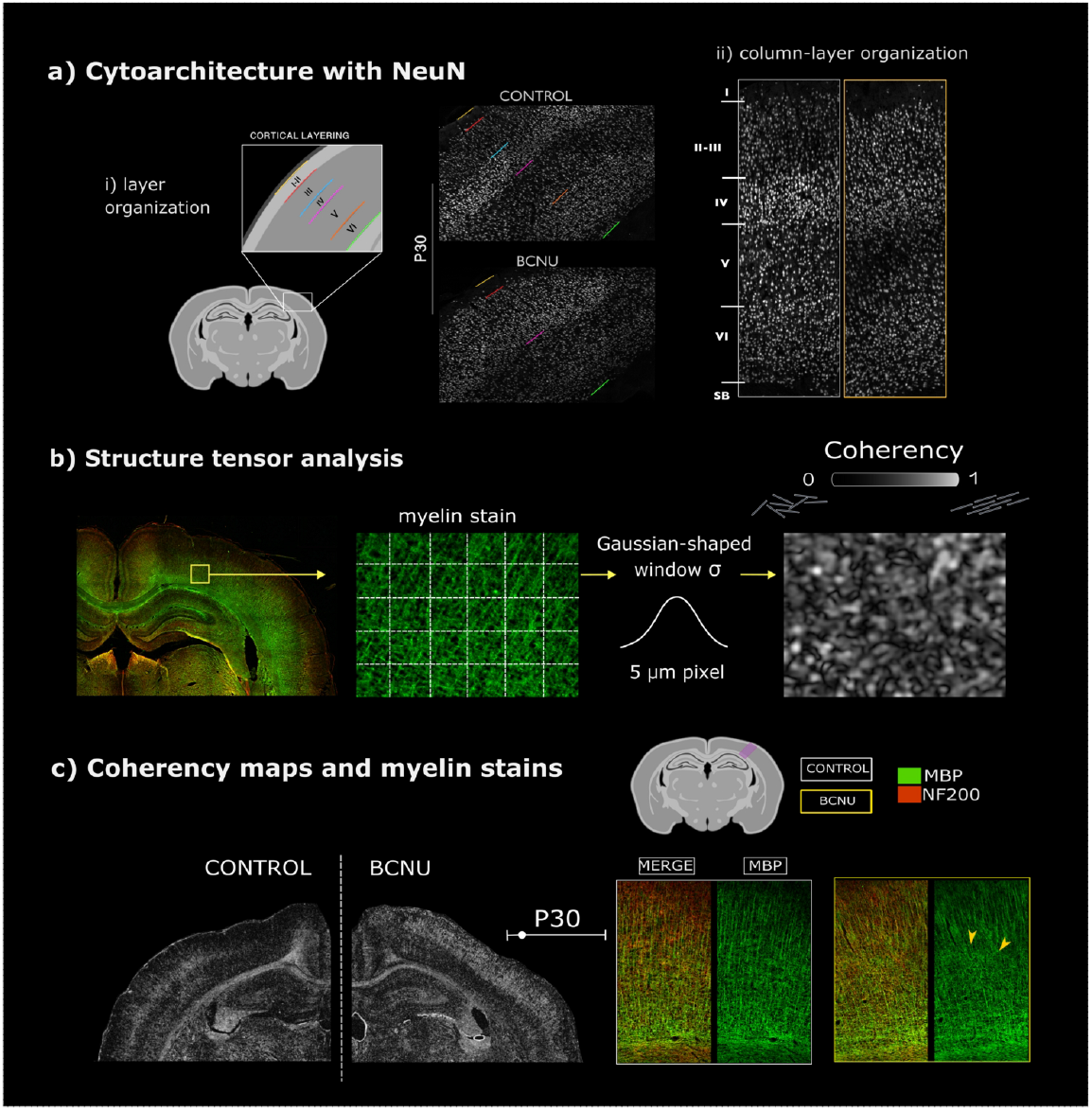
Representative stainings for cyto- and myeloarchitecture and structure tensor at P30. a) Coronal sections at the level of the dorsal hippocampus stained with NeuN revealed alterations in their layering organization. a) Layer organization was delineated by color-coded lines as indicated in (i). Close-up of the somatosensory cortex in BCNU-treated animals shows blurred distinction between cortical layers III-IV (cyan line) and V-VI (orange line), in contrast to control animals. A closer examination of columnar-layer organization (ii) evidences a clear layer differentiation in control rats (white rectangle) accompanied by radial neuronal orientation. BCNU-treated rats (yellow rectangle) displayed a series of abnormalities including disorganized vertical orientation, decreased cellular density across layer V, and slightly enlarged cell bodies. b) MBP stains (in green) were used to estimate coherency maps through structure tensor analysis. The resulting maps are visualized using a gray scale: brighter regions indicate higher fiber organization (i.e., coherency), and darker regions represent lower textural coherence. BCNU-treated rats exhibited decreased coherence between the white and gray matter boundaries, particularly at P30 (arrows). Myelin-stained sections from the somatosensory cortex (right panel) revealed disrupted myelin fiber arrangements (yellow arrows) in layers VI and V of BCNU-treated rats (yellow boxes) compared to control animals (white boxes).

Cortical disarray not only impacted the layer organization but also affected columnar assembling, as seen in Fig. 7a-ii. In control animals, chains of neurons followed a radial orientation from the deepest layer (VI) to the superficial layer II over time after birth. On the contrary, BCNU-treated rats showed not only a disorganized vertical orientation, but also a qualitative reduction in cell density, particularly in layer V, and an increased cell body size, with the most pronounced effects observed at P60.

#### 3.3.2 Myeloarchitecture and structure tensor analysis

Our examination of myelin-labeled samples provided valuable insights into the microstructural organization of intra-cortical fibers. Using the structure tensor analysis workflow, we investigated the fiber structure in both groups by computing the coherency maps **(Fig**. 7b), and staining for myelin (green) and neurofilament (red) to elucidate their potential impact on cortical diffusivities. Coherency maps (Fig. 7c; left panel) derived from myelin staining depicted consistent anisotropic patterns in cortical layers IV and VI in control animals, with slight age-dependent variation in intensity. However, in the BCNU-treated sample at P30, a significant cortical area displayed abnormal coherency spanning from layer IV to I, accompanied by a loss of orientation in the deeper layers V and VI. Corresponding myelinated fibers taken from the somatosensory cortex exhibited radial and intracortical fiber disarrangement in the BCNU-treated rat. While pronounced changes in primary orientation were evident at P30, similar alterations persisted in subsequent developmental stages, albeit at a lesser degree **(Supplementary Fig. S6)**.

## 4 Discussion

The cerebral neocortex comprises a highly organized arrangement of cells and fibers, rendering it a complex structure that supports high-level functions. While histology has yielded profound insights into cortical microstructure, challenges persist in achieving microstructural specificity through *in vivo* MRI. This limitation not only poses hurdles to scientific inquiry but also hinders clinical applications aimed to detect cortical abnormalities at the mesoscale level. In this study, we provide evidence supporting the use of advanced analysis of voxel-wise and per-bundle diffusion metrics to detect cortical orientational microstructural components in a rat model of cortical dysplasia.

Diffusion-weighted MRI opened an important avenue for non-invasivc tissue exploration beyond macrostructural features. While most diffusion-weighted MRI methods have focused on studying white matter bundles and structural brain connectivity, there is a growing trend toward unraveling the complexity of the cortex. The neuropil (i.e., the space between neurons and glial cells, including axons, dendrites, and microvasculature) [71], has such an intricate organization that it-is expected to show little diffusion anisotropy through conventional diffusion-weighted MRI methods (Kroenke, 2018). However, despite lacking large-caliber axons like those in white matter, the histological configuration of the neocortex is not isotropic. In contrast, the dense radial and perpendicular arrangement of neurons and myelinated fibers could potentially contribute to the anisotropic diffusion of the cortex [11, 67, 68]. For instance, cortical regions with well-defined columnar and layer boundaries, such as the motor and somatosensory cortices, have been parcellated by fitting DTI [8, 34, 53, 57]. Previous studies have also shown a strong radial diffusion component inside the cortex with variations of the tangential orientation across cortical areas [5, 49, 53]. These features are lost in focal cortical dysplasia, as evidenced by high resolution DTI of excised brain tissue of patients with epilepsy that does not respond to pharmacological treatment [31].

Our work shows that methods exploiting non-single Gaussian diffusivity are more sensitive in capturing generalized cortical microstructural abnormalities, while per-bundle analysis of the diffusion signal can disentangle the contribution of fibers and cellular arrangements that run radial or tangential to the cortical surface in the rodent brain, surpassing the limitations inherent in single-tensor techniques. We show that several diffusion metrics are able to detect abnormalities of cortical microstructure even in the absence of gross morphological alterations. This becomes particularly relevant for the detection of very subtle cortical alterations that can nonetheless cause severe neurological disorders, such as medically-refractory epilepsy due to focal cortical dysplasia [14]. To prove the validity of our methodological approach, we used an animal model of FCD which, through the administration of carmustine (BCNU) during a critical stage of cortical development, causes histological disarrangement of the cortex with features akin to those seen in FCD Type II [15]. While other animal models of FCD better reflect the electrophysiological properties of these malformations of cortical development [51, 60, 80], the BCNU model used here serves as a valid proxy for their histological abnormalities and a good fit for their detection with diffusion-weighted MRI. Our longitudinal design further allowed us to evaluate the trajectories of these changes over time. In previous studies, T2-weighted images have shown that BCNU-treated rodents displayed macrostructural abnormalities such as ventricular enlargement and hippocampal atrophy at P60 [40, 54, 55], similar to those depicted in Fig. 2. Here, ventricular enlargement and atrophy became apparent from P60 onwards, yet-diffusion abnormalities were evident-at P30.

While microscopic developmental changes such as fiber organization, maturation of oligodendrocyte and neuronal migration toward the cortical surface [26, 82]-often go unnoticed in anatomical images, in this study we proved that diffusion-weighted MRI is sensitive to microstructural changes induced by cortical dysplasia. For example, mean kurtosis-which reflects the heterogeneity of an environment where water molecules experience physical barriers-revealed differences between groups at P30 (Fig. 5-ii). Diffusion kurtosis was higher in control animals, likely reflecting a more complex and compact laminar structure compared to the BCNU group. When examining the group-average kurtosis, regional differences became more apparent. As shown in Fig. 3, the control group exhibited peak kurtosis in the deepest cortical layers with noticeable gradient toward the more superficial layers, likely reflecting a switch from compact to less densely packed medium. In contrast, the BCNU group showed a more homogenous kurtosis across cortical depth, meaning overall lower restriction and heterogeneity among layers without a clear transition referring to a unique layer composition.

DKI proved useful for detecting overall changes in microstructural restriction paralleled by histological features. NeuN staining provided clear evidence of disrupted laminar architecture, showing blurred boundaries between cortical layers and, to some extent, disrupted columnar organization. The loss of horizontal delineation likely reduces restriction in the vertical direction, allowing water to diffuse more freely in that direction. This is consistent with the observed reduction of AK—and, to a lesser extent, RK—in BCNU rats relative to controls (Fig. 5-ii), which in turn explains the overall MK trend driven mostly by RK.

Voxel-wise summary metrics derived from DTI, DKI and NODDI are insufficient to capture complex orientational information, particularly the organization of neocortical myeloarchitecture into radial and tangential fibers. Per-bundle diffusion metrics derived from CSD and MRDS showed distinct abnormalities in BCNU-treated rats. AFD_rad_ was the most sensitive metric at P30, with increased values throughout the cortex. Notably, FA_rad_ and FA_tan_ showed reductions in the deep layers at the same age. This apparent discrepancy may be explained by a paucity of tangential fibers in experimental animals, as evidenced by their abnormally high coherency index derived from MBP labeling (**Fig**. 7). In regions of fiber crossings with asymmetric histological characteristics, AFD can be artificially increased in the dominant fiber bundle when a fixed response function is used [69]. Increased fiber dispersion can also cause increased AFD at the expense of reduced FA, but ODI maps derived from NODDI (**Figs**. 3 and 5) argue against this interpretation.

The evolution over time of diffusion metrics and between-group differences showed a complex pattern. Our results show that P30 represents a development inflection point in this animal model of cortical dysplasia. Around this time, animals show rapidly diverging microstructural trajectories, which later plateau into a more stable—though still abnormal—architectural pattern. The conjunction of diffusion abnormalities observed at P30 suggest that BCNU-treated rats present a disruption in their cortical maturation during critical developmental periods, and potentially reach a plateau in later postnatal ages. Such observations were consistent-with our histological findings, where radial and tangential fiber disorganization was detected in myelin-stained samples in BCNU-treated rats (**Fig**. 7), but-tended to normalize over time **(Supplementary Fig. S6)**.

There are important considerations about our study. First, it involves the direct application of diffusion methods originally designed for the analysis of white matter to study the neocortex. While CSD and MRDS have demonstrated robustness in estimating tensors in areas with crossing fibers [21, 69], their reliability in gray matter remains a subject of inquiry. However, our prior investigation [77] supports evidence that MRDS can detect changes of cortical organization at the early stages of development. Similarly, per-bundle analysis of AFD (i.e., fixel-bascd) was also originally conceived for analysis of white matter bundles providing robustness to fiber crossings. Our work is in line with previous reports that analyze high-resolution diffusion-weighted MRI obtained *ex vivo* [49] showing the applicability of fixel-wise analysis within the cortical mantle. Second, this study undertook a longitudinal analysis involving the sacrifice of four animals (n=2 Control; n=2 BCNU) at each temporal point. This approach introduces two implications: 1) the reduced number of samples at each point-that could impact the statistical analysis, and 2) the lack of one-to-one correlation between total diffusion-weighted MRI metrics and histology. Third, while our images possess high spatial resolution in-plane, slice thickness is relatively large (1 mm), thereby introducing partial volume effects in our diffusion-weighted MRI estimations. Next, while MRDS can fit-up to three tensors, in this analysis we only included the most radial and most tangential tensors with respect to the grid-line. Similar approaches were taken with CSD as we constrained the fixel calculation per voxel to three, and applied the same condition to select the radial and tangential representative fixels. These tangential tensors/fixel (representing fibers running tangential to the pial surface) could have a medial-to-lateral orientation, or run orthogonal to the imaging plane. The distinction between these two forms of intra-cortical diffusivities (i.e., intracortical fibers) was not attempted due to the aforementioned thick slices. Finally, while the methods described here can be applied to the human condition, the spatial resolution afforded clinical scanners is much lower than what can be achieved with pre-clinical scanners. On the other hand, the possibility of diffusion weighted MRI to discern structural features within voxel, advances in high-performance gradients [29], and the advent of super-resolution techniques [22, 52] will propel the adoption of diffusion-weighted MRI for the study of neocortical features.

Our study offers compelling evidence that DKI and per-bundle MRDS and CSD can discern microstructural abnormalities associated with abnormal organization of cortical layers and columns. Moreover, it proves useful for separating and identifying the orientational cortical components of cyto- and myelo-architecture. While this study is preclinical in nature, our findings underscore the potential of utilizing dMRI and advanced multi-tensor methods for clinical applications where neocortical histopathology is suspected.

## Acknowledgements and Funding

We thank Mirelta Regalado, Ana Aquiles, Juan Ortiz-Retana, Nydia Hernández-Ríos, Ericka de los Ríos, Lourdes Palma, Gema Martínez, and Leopoldo Gonzálcz-Santos for their technical assistance. We also thank the personnel at the Animal Facility: Martin Garcia, Alejandra Castilla, Maria A. Carbajo, and Maria E. Ramos. This research was funded by CONAHCYT (CF-218-2023 for LC) and UNAM-DGAPA-PAPIIT (IN2044720 and IN213423 for LC; IA200621 and IN211326 for HL-M. MRI preclinical scanner facilities were provided by the National Laboratory of Magnetic Resonance (LANIREM), which receives support from CONAHCYT and UNAM. Computing infrastructure was partially provided by the National Laboratory for Advanced Scientific Visualization (LAVIS). Paulina J. Villaseñor received a scholarship from CONAHCYT (798166). AR-M was partially supported by CONAHCYT.

## Author contributions statement

PJV, HL-M, AR-M, and LC contributed to the conception, methodology, and design of the study. HL-M and PJV performed experiments and imaging acquisition. PJV performed diffusion-weighted MRI statistical analysis, histological procedures, and structure tensor analysis. RC-L provided the MRDS code and supervised the analysis of diffusion. PJV and LC wrote, reviewed, and edited the first manuscript. All authors reviewed the manuscript and approved the submitted version.

## Code availability

Vertex-wise statistical workflow using the LMM and cluster-wise is available in this repository: https://github.com/paulinajv/Vertex-wise-LMM-analysis Pipeline to generate the 2D grid-lines is available in this repository: https://github.com/lconcha/Displasias. MRDS code is available under request

## Supplementary information

**Supplementary Figure S1:**
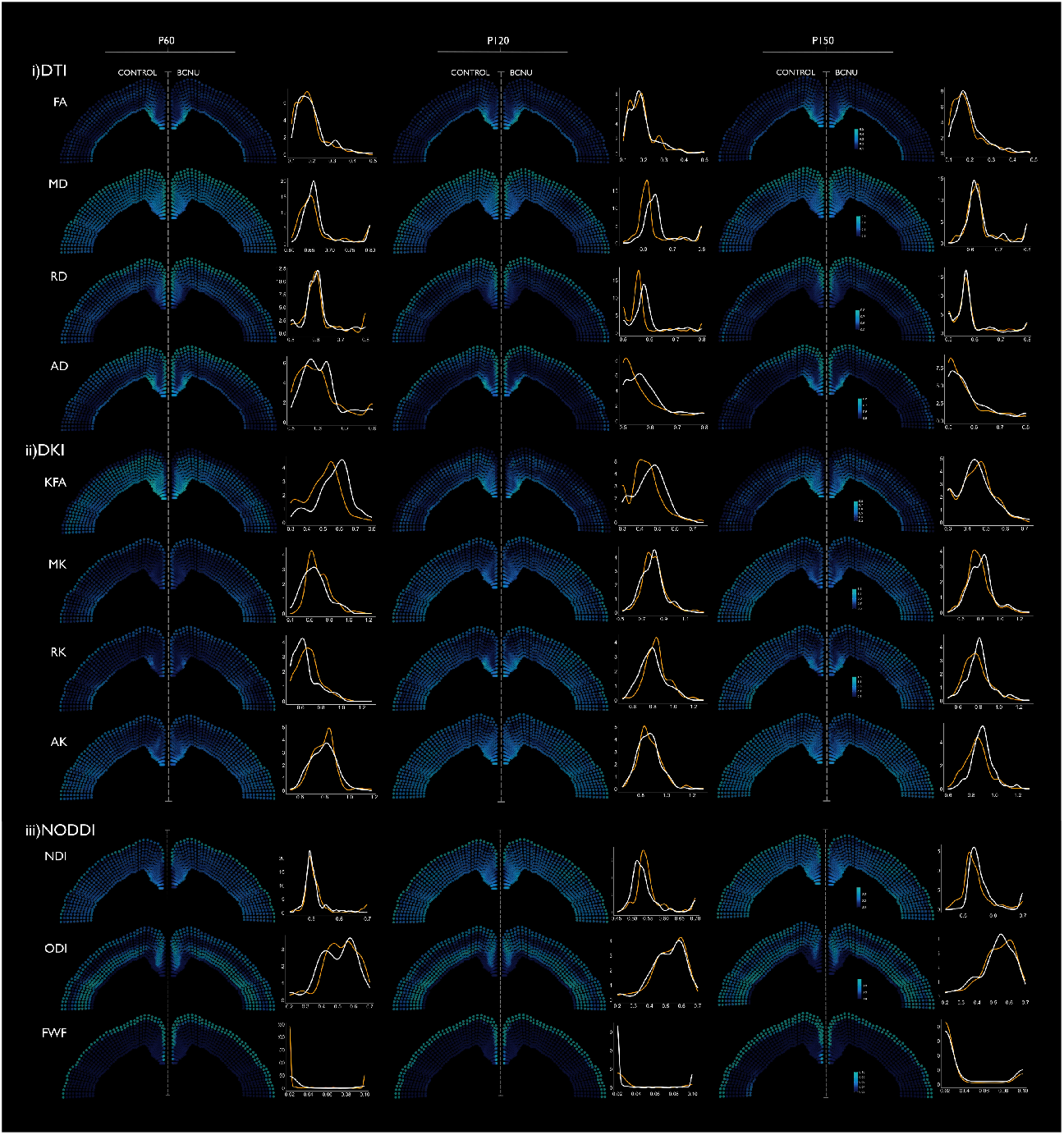
Longitudinal vertex-wise means for diffusion MRI scalars with corresponding density distributions. For each vertex along the grid-line, the mean values of i) DTI and ii) DKI and iii) NODDI metrics were calculated separately for the control (shown as left hemisphere) and BCNU (shown as right hemisphere) groups at P60, P120 and P150 (see **Fig**. 3 for P30). Mean values are displayed in the same color scale for both groups to facilitate comparison. Density curves describe the frequency distribution rates of each diffusion metric across all the grid-line vertices for both groups (BCNU in yellow; control in white)

**Supplementary Figure S2:**
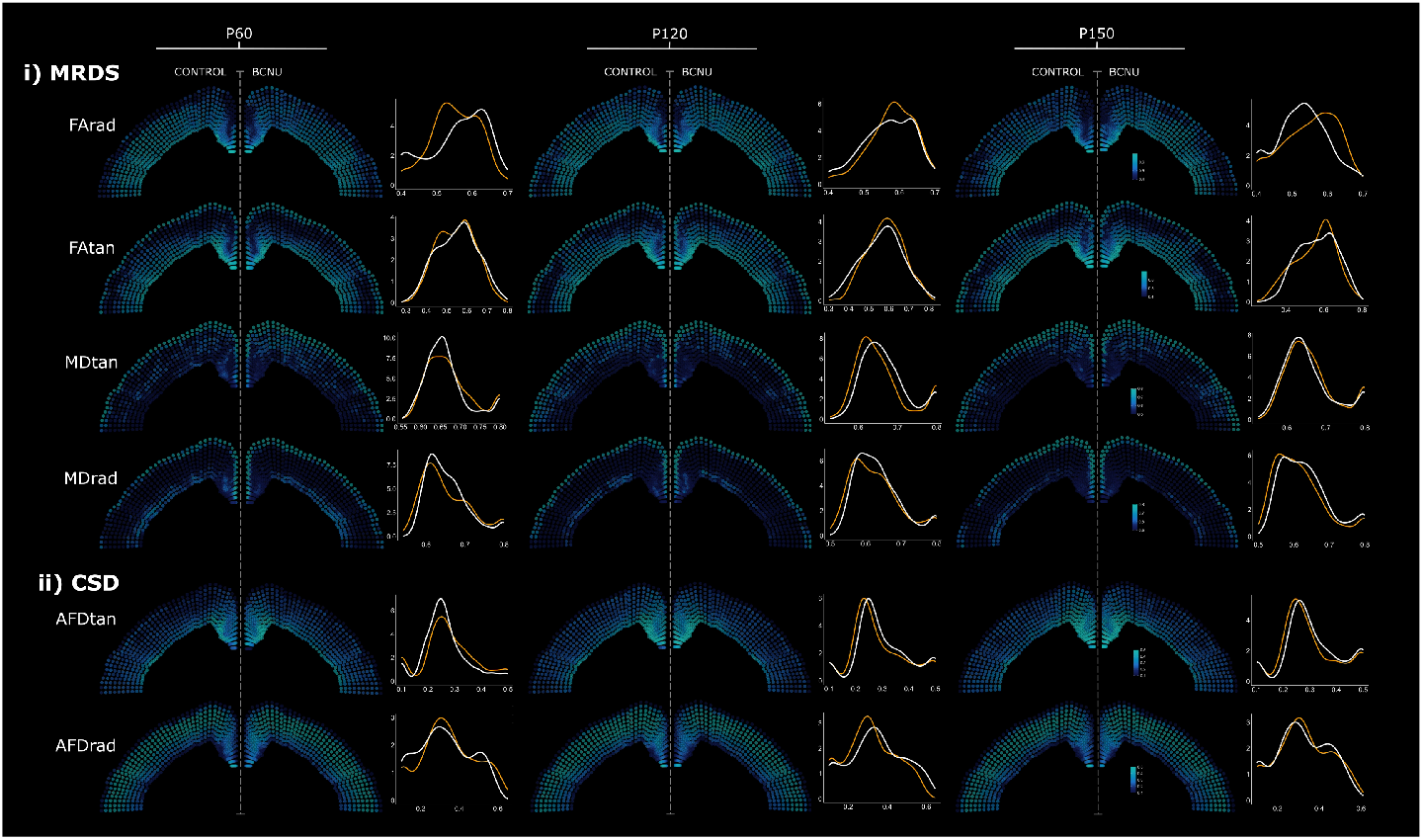
Longitudinal vertex-wise means for diffusion MRI per-bundle with corresponding density distributions. For each vertex along the grid-line, the mean value of i) MRDS, and ii) CSD metrics was calculated separately for the control (shown as left hemisphere) and BCNU (displayed as right hemisphere) groups at P60, P120 and P150 (see **Fig**. 4 for P30). Display conventions as in **Supplementary Fig**. SI.

**Supplementary Figure S3:**
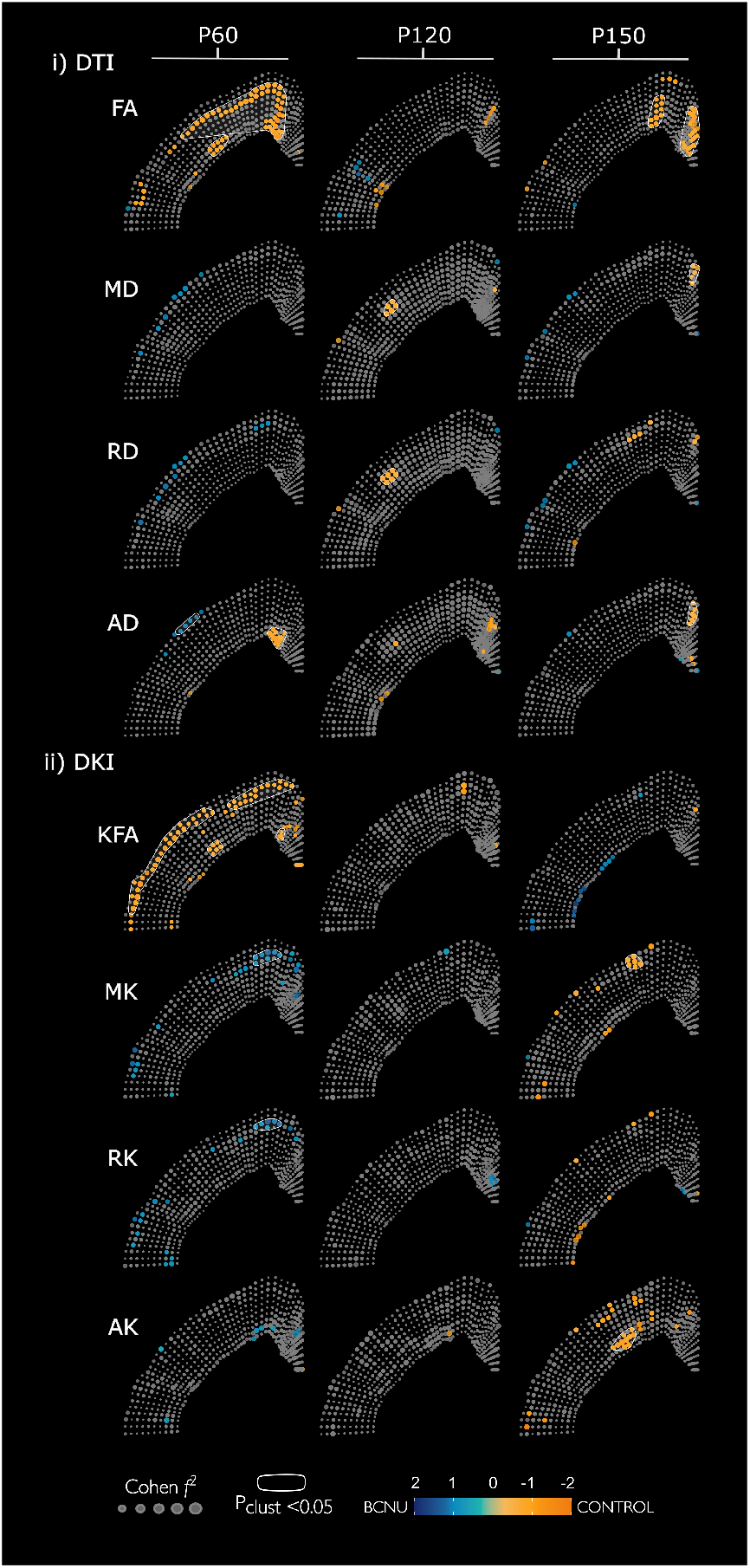
Vertex-wise DTI and DKI analysis of the cortex at P60, P120 and P150. Significant point-wise post hoc between-group differences (p<0.05) are represented in colors (cool colors for reductions and warm colors for increases in experimental animals with respect to control animals); non-significant in gray, with absolute effect size represented as marker size. See **Fig**. 5 for data related to P30.

**Supplementary Figure S4:**
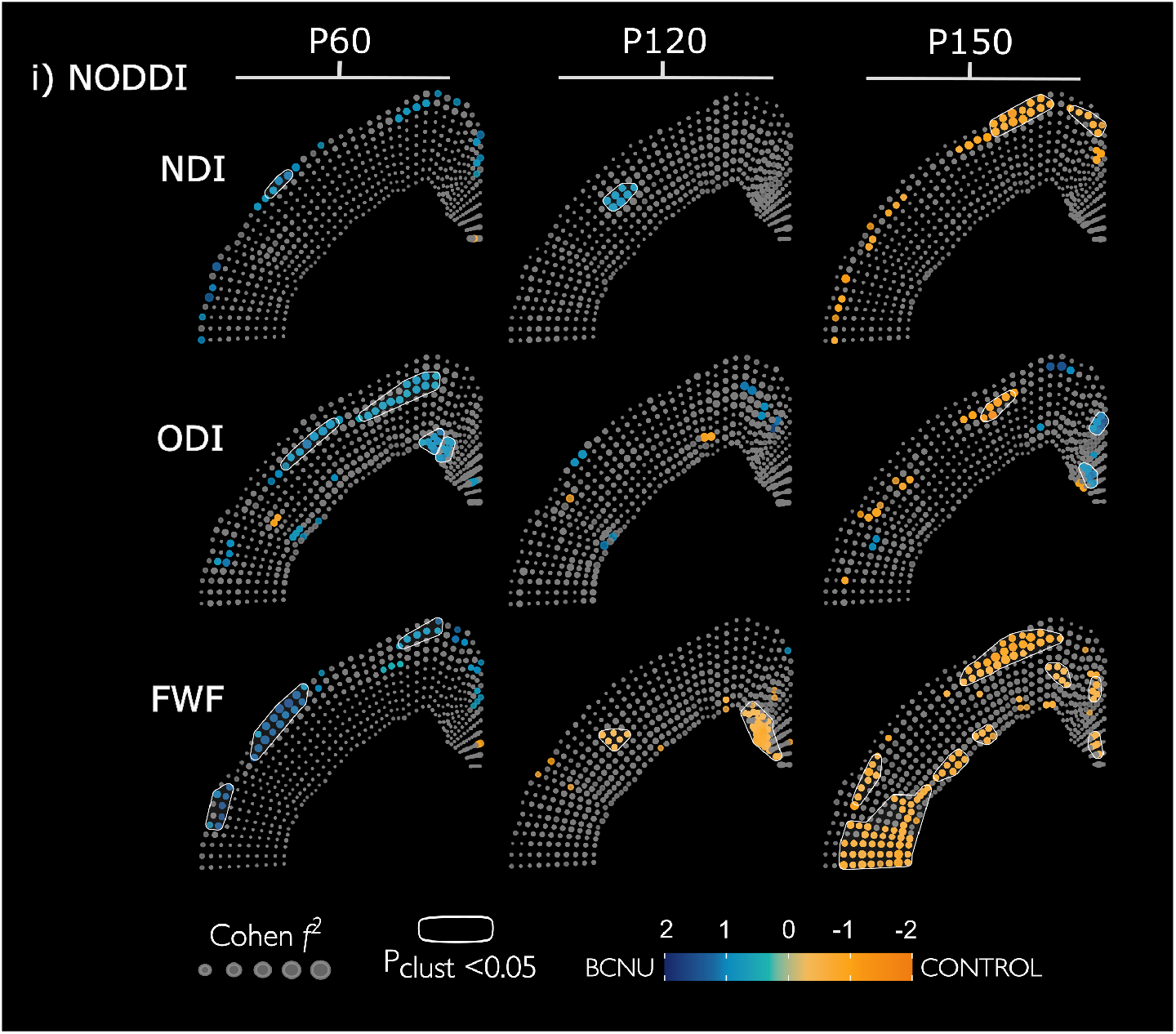
Vertex-wise NODDI analysis of the cortex at P60, P120 and P150. Plotting conventions as in **Supplementary Fig**. S1. See **Fig**. 5 for data related to P30.

**Supplementary Figure S5:**
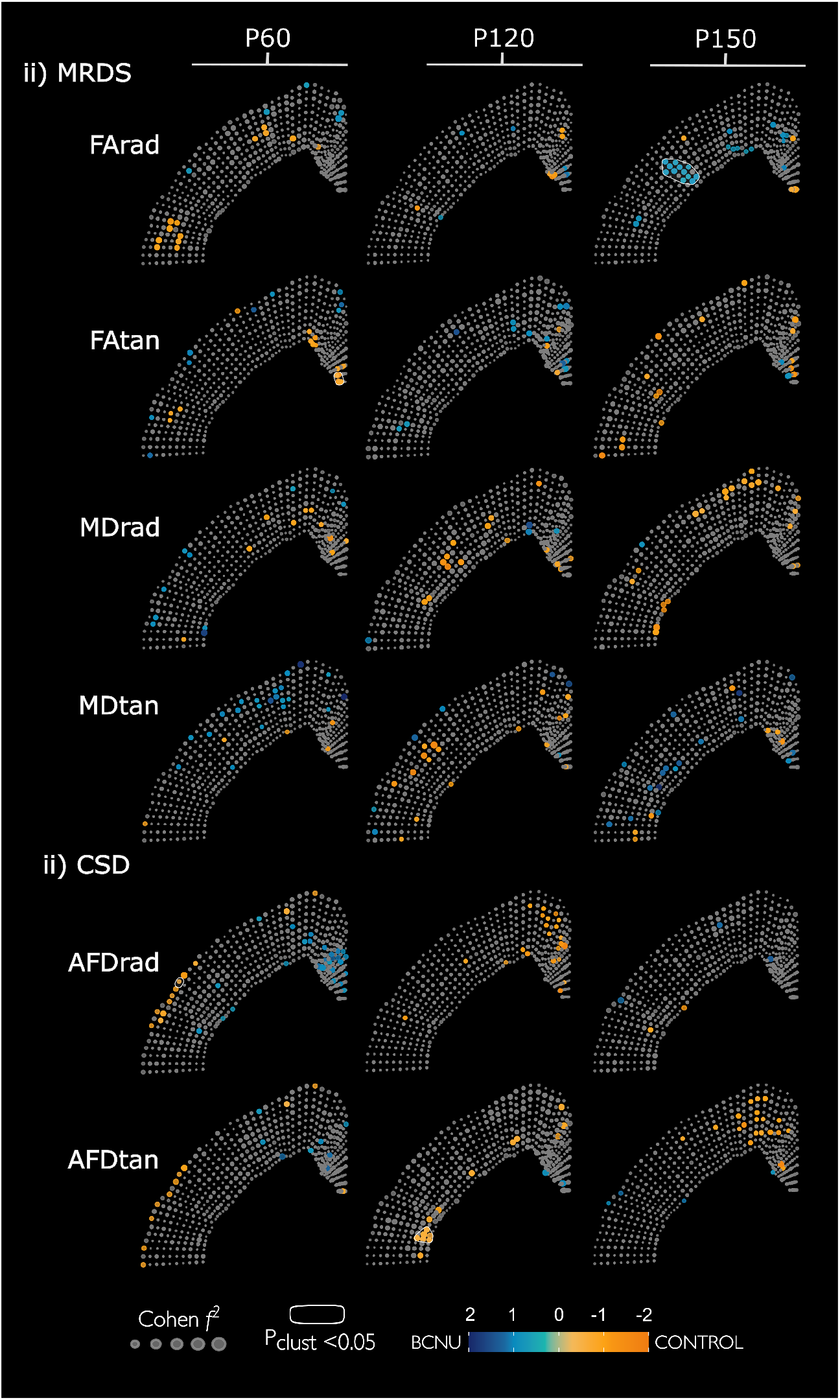
Per-bundle vertex-wise analyses with a multi-tensor fit (MRDS, i), and CSD (ii) at P60, P120 and P150. Plotting conventions as in **Supplementary Fig**. S1. See **Fig** 5 for data related to P30.

**Supplementary Figure S6:**
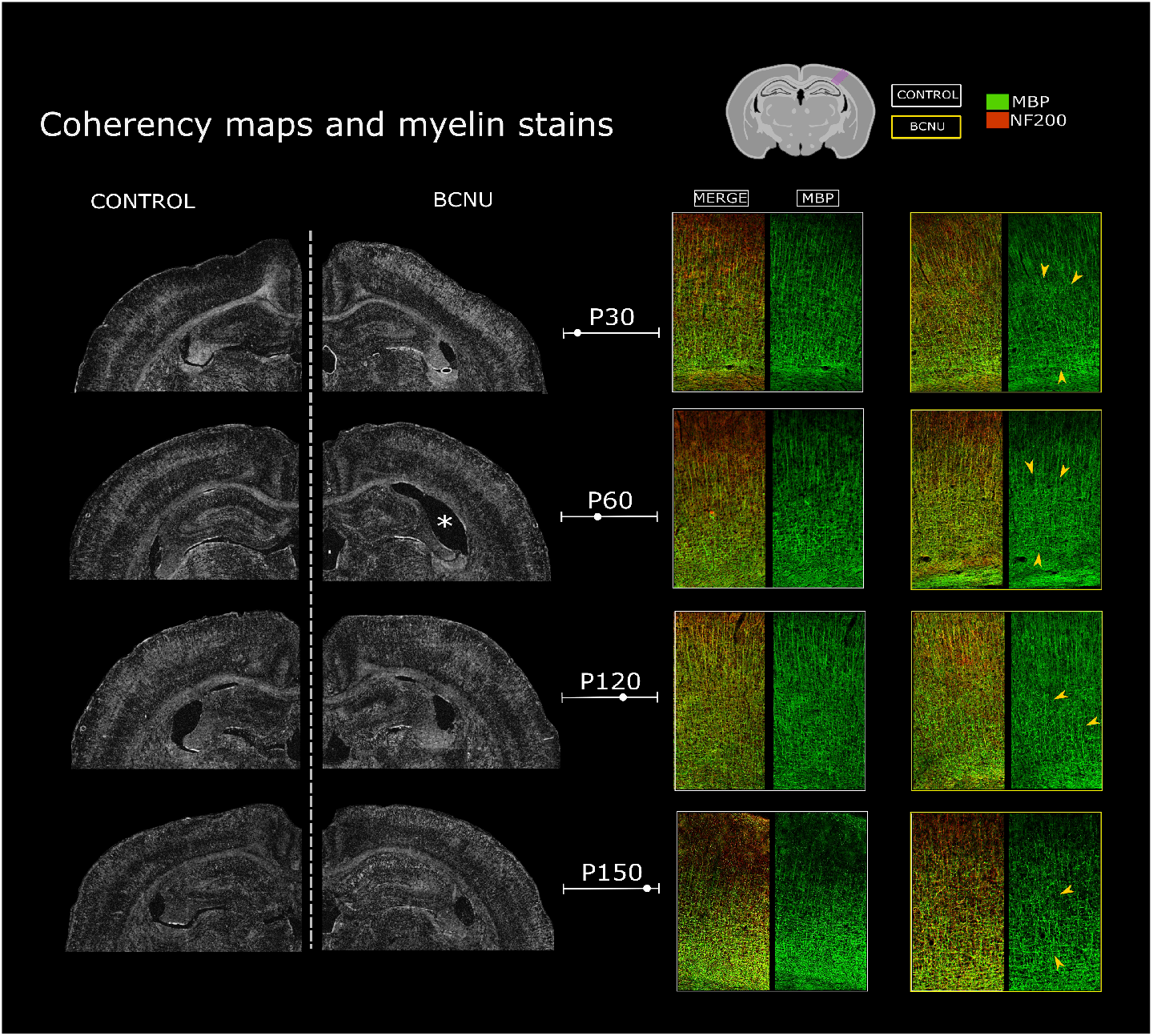
Longitudinal Structure tensor analysis derived from MBP immunofluorescence. Coherency maps indicate the level of organization of MBP+ fibers within a local window around each pixel (Gaussian kernel of 5 μm). Coherency ranges from zero (black, completely incoherent orientations) to one (white, a main orientation is clear). Asterisk shows an enlarged ventricle. High magnifications of an area indicated by the rectangle in the atlas are shown on the right. Yellow arrowheads point at disorganized myelinated fibers.

## Notes

### Competing Interest Statement

The authors have declared no competing interest.

### Summary of Updates

Affiliations for some authors were incorrect. This was fixed.

